# Differential and temporally dynamic involvement of primate amygdala nuclei in face animacy and reward information processing

**DOI:** 10.1101/2024.01.16.575972

**Authors:** Koji Kuraoka, Kae Nakamura

## Abstract

Decision-making is influenced by both expected reward and social factors, such as who offered the outcomes. Thus, although a reward might originally be independent from social factors, the two elements are closely related. However, whether and how they are processed separately or conjointly remains unclear. Here, we show that neurons in distinct sub-nuclei of the amygdala encode expected reward and face animacy, which is a vital aspect of face perception. Although these encoding processes are distinct, they rely on partially shared neuronal circuits with characteristic temporal dynamics.

Two male macaque monkeys made saccades under different social and reward contexts, created by presenting facial images with independent attributes: animacy (a monkey or cartoon face) and associated reward (large or small). The stimulus image was presented twice per trial: during the initial stimulus encoding (S1) and before saccades were made (S2). A longer gaze duration for eye region of the monkey versus cartoon images indicated more robust social engagement for realistic faces. During S1, a similar number of lateral nucleus neurons encoded either animacy only with a monkey-image preference, reward only with a large-reward preference, or both. Conversely, neurons in the basal and central nuclei primarily encoded reward, preferring large-versus small-reward associated face images. The reward-dependent modulation was continuous after S1, but was more conspicuous during S1 in the basal nucleus and during both S1 and S2 in the central nucleus. This anatomically- and temporally-specific encoding in the amygdala may underlie the computation and integration of face animacy and reward information.

**Significance Statement:** Reward and social information are closely related but originally independent, as both influence our decision-making. The amygdala has been associated with both reward and social information coding. However, whether and how they are processed separately or conjointly by individual neurons in the amygdala remains unclear.

We found that neurons in the lateral and basal nuclei encoded face animacy, which is an important aspect of social information, and reward, respectively, during sensory processing. Neurons in the central nucleus encoded reward information during the execution phase. This provides new clarity regarding the mechanisms of separate or integrated social and reward information processing within the amygdala.

## Introduction

Decision-making is influenced by both an expected reward and social aspects, such as who offered the outcomes. For example, at a restaurant, we may perceive the same food differently depending on whether it is served by a robot or human waiter. Thus, although reward and social information might originally be independent, they are closely related.

Among the many brain areas implicated in reward coding, the amygdala plays a key role in value evaluation of sensory events (Paton et al., 2006; Belova et al., 2007; Salzman et al., 2007; Belova et al., 2008; Morrison and Salzman, 2009; Shabel and Janak, 2009; Janak and Tye, 2015; Namburi et al., 2015; Grabenhorst et al., 2016; O’Neill et al., 2018; Pignatelli and Beyeler, 2019).

The amygdala has also been associated with social cognition (Adolphs, 2010). Bilateral damage to the amygdala can impair perception regarding the intensity of fearful facial expressions (Adolphs et al., 1994). Single neurons in the primate amygdala encode integrated information about facial identity and expressions (Kuraoka and Nakamura, 2006; Gothard et al., 2007) and eye contact (Mosher et al., 2014). Amygdala neurons also signal the value of rewards delivered to the self and to others (Chang et al., 2015), as well as social decisions (Grabenhorst and Schultz, 2021).

Detecting animacy, a property used to distinguish animate from inanimate entities, such as humans from robots, is a critical skill for successful social interactions (Shultz et al., 2015). Human imaging studies have shown that the amygdala responds more strongly to human faces or animal images than to robotic faces or abiotic objects, suggesting its involvement in animacy perception (Yang et al., 2012; Coker-Appiah et al., 2013; White et al., 2014). Consistently, patients with amygdala lesions lack the ability to anthropomorphize objects (Heberlein and Adolphs, 2004; Waytz et al., 2019), suggesting a critical role of the amygdala in animacy. However, whether and how individual neurons in the amygdala encode reward and animacy information is not clear.

The lateral (La), basal (Ba), accessory basal (AB), medial (Me), and central (Ce) sub-nuclei of the amygdala receive projections from distinct brain areas (Freese and Amaral, 2009) in the reward system (Schultz, 2016) and social networks (Kennedy and Adolphs, 2012). Previous studies have examined the distribution of neurons that process reward or social information in the sub-nuclei. One tested the selectivity of single amygdala neurons for social nature (monkey versus object), reward (three different juice amounts), task event (fixation, stimulus onset, stimulus offset), and stimulus-unique features (face or eyes), and reported that neurons with multiple selectivity were distributed among all major sub-nuclei of the amygdala (Putnam and Gothard, 2019). Other groups reported shared coding for reward and more complex social information such as social hierarchy within conspecific face images (Munuera et al., 2018) and eye gaze direction, which is an important aspect of social information (Pryluk et al., 2020), in neurons in the primate basolateral complex. However, the detailed distribution of neurons that encode complex social and/or reward information remains controversial.

The amygdala has also been implicated in different aspects of task sequences, such as stimulus-reward associations (Spiegler and Mishkin, 1981) and decision-making (Grabenhorst et al., 2012). Thus, social and reward information processing may vary according to sensory information encoding and motor execution.

In this report, we examined 1) whether distinct amygdala regions compute face animacy (i.e., complex social information) or reward information separately or conjointly, and 2) whether face animacy and reward coding vary according to the stage in a concurrent cognitive process (sensory information encoding or motor execution). To this end, we developed a behavioral paradigm in which animals made saccades under different animacy conditions (viewing a real monkey face or a cartoon face) and reward contexts (predicting large or small rewards). We measured animacy and reward coding during initial stimulus-encoding and later execution processes. We found anatomically- and temporally-specific encodings of face animacy and reward in distinct sub-nuclei of the primate amygdala.

## Materials and methods

### General

We used two hemispheres of two male Japanese monkeys (*Macaca fuscata,* laboratory designations: Animal P; 9 years old, 9 kg, and C; 7 years old, 11 kg). All experimental procedures were performed in accordance with the National Institutes of Health Guidelines for the Care and Use of Laboratory Animals and were approved by the Institutional Animal Care and Use Committee at Kansai Medical University.

Each animal underwent implantation of a head post to allow maintenance of a stable head position. Eye position was monitored using an infrared video-tracking system with a time resolution of 500 Hz and a spatial resolution of 0.25–0.5 deg (iView X 2.8; SensoMotoric Instruments, Boston, MA, USA). All experiments were performed in a dark soundproof room, where the macaques sat in a primate chair and faced a 24-inch monitor (ProLite B2403WS; Iiyama, Tokyo, Japan) positioned 38 cm from their eyes. All aspects of the behavioral experiment, including stimuli presentation, monitoring of eye movements and neuronal activity, and outcome delivery were controlled by a real-time experimentation data acquisition system (Tempo; Reflective Computing, WA, USA).

### Behavioral tasks

Face stimuli are well-known animate agents (Shultz et al., 2015). Comparing real and artificial faces is an effective way to examine face animacy perception (Gobbini et al., 2011; Balas and Koldewyn, 2013; Wang et al., 2020). Accordingly, we used a set of eight face stimuli, composed of four different real monkey faces and four cartoon faces shown in different colors (Fig. 1A). All face stimuli had a uniform mean luminance and size (number of pixels). Two monkey faces (M1L, M2L, Fig.1A) and two cartoon faces (C1L, C2L) were consistently associated with a large reward, whereas the other two monkey faces (M3S, M4S) and two cartoon faces (C3S, C4S) were consistently associated with a small reward. Both animals learned to associate each face with the amount of the expected reward through extensive training.

**Figure 1.**
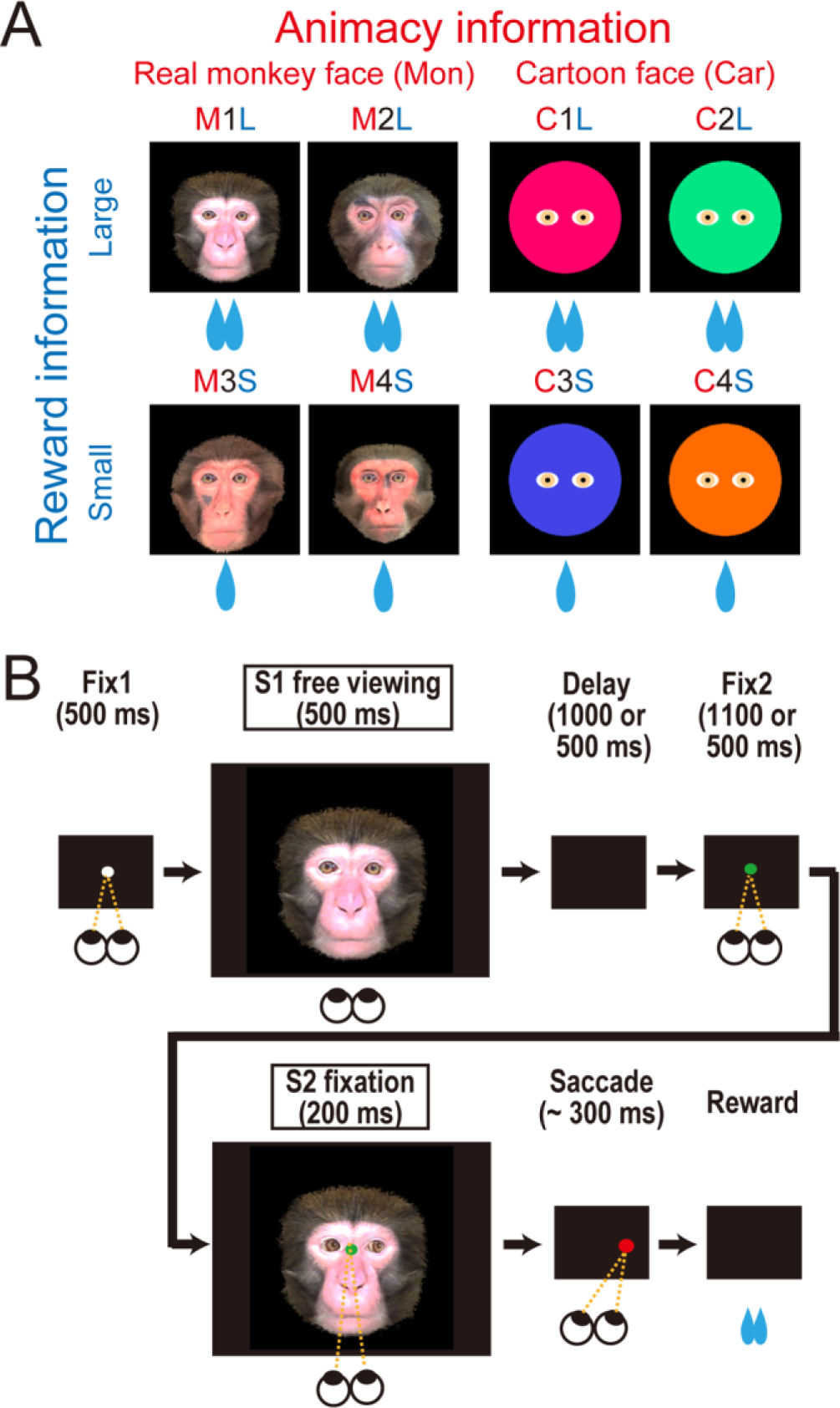
Experimental paradigm. **A**. Visual stimuli with two attributes: degree of animacy, i.e., monkey (Mon) or cartoon (Car), and reward size, i.e., large or small. **B**. Visually-guided saccade task under different animacy and reward contexts. After fixation on fixation point 1 (Fix1) for 500 ms, 1 of 8 visual stimuli (A) was briefly presented (S1), followed by a delay (1000 or 500 ms) and the presentation of fixation point 2 (Fix2). After fixation on Fix2, we presented a visual stimulus (S2), which was the same as S1 except for the gaze direction (see Extended Figure 1-1), followed by a target. The animal was required to make a visually guided saccade towards the target to receive a large or small reward, as indicated by S1 and S2 (M1L, M2L, C1L, and C2L for a large reward; M3S, M4S, C3S, and C4S for a small reward).

We presented the stimuli twice while each animal performed a visually-guided saccade to obtain a liquid reward (Fig. 1B). After fixating on the first central fixation point (Fix1) for 500 ms, the first face stimulus (S1) was presented for 500 ms. The animals were allowed to scan the image freely. After a delay with a blank screen (1000 ms for Animal P; 500 ms for Animal C), the second central fixation point (Fix2) appeared. After fixating on Fix2 for 1100 ms (Animal P) or 500 ms (Animal C), the second face stimulus (S2) was presented briefly (200 ms), followed by the presentation of a target dot on the left or right, 8 deg from the center. The animals were then required to make a saccade towards the target to obtain a liquid reward.

The first face stimulus (S1) always looked forward. The second face stimulus (S2) was identical to S1 except that the gaze direction of the face was either toward the left or right of the screen. As shown in Supplementary Figure 1-1, half of the faces (M1L, M3S, C1L, C3S in Fig. 1A) always looked toward the future target (‘Congruent faces’), while the other half (M2L, M4S, C2L, C4S in Fig. 1A) always looked away from the future target (‘Incongruent faces’). Thus, visual stimulus S1 carried information about (1) the degree of animacy (real monkey or cartoon face), (2) the expected reward size (large or small), and (3) the congruency of the gaze direction between S1, S2, and the saccade target. The present manuscript did not address (3) because the behavioral and neural effects were small.

To evaluate each animal’s preference for the stimuli, we also conducted a stimulus-assessment task (Supplementary Fig. 2-1). In this task, the subjects were forced to choose one of two face stimuli that were simultaneously presented as S1. Specifically, they had to choose between a monkey and cartoon face or between a large- and small-reward associated face. The chosen stimulus was presented in the following S2 period. The stimulus preference was evaluated according to the proportion of each stimulus choice.

### Recording neuronal activity and localization of the amygdala nuclei

We recorded extracellular activity from single neurons in the amygdala subnuclei using tungsten electrodes (diameter, 0.25 mm; impedance, 0.5–2 M Ω at 1 kHz; Frederick Haer). The signal was amplified with a bandpass filter (300 Hz–8 kHz; MCP-Plus 8; Alpha Omega LT) and collected at 1 kHz. The single-neuron activity was isolated and converted into pulses via a template-matching protocol (20 kHz for waveform matching and spike sampling; Power1401-3A; CED).

To enable recording, we implanted a circular plastic recording chamber (Crist Instruments, MD, USA) at 20 mms AP and 11 mms lateral with a 10-degree lateral tilt. The location of the amygdala was verified by overlaying penetration record maps on magnetic resonance images (MRI) (for Animal P, 1.5 T, SIGNA, General Electric Company, CT, USA, for Animal C, 0.3 T, AIRIS, Hitachi, Tokyo, Japan). MRI data were obtained with the recording chamber filled with gadolinium, which clarified the relative location of the recording chamber and the underlying brain structure. The electrode was inserted into the brain through a stainless-steel guide tube (0.9 mm in diameter) that was fixed to a grid system. The guide tube was inserted through the dura to a depth of ∼5 mm above the amygdala, as estimated using the MRI data. The electrode was advanced using a hydraulic microdrive (Narishige, Tokyo, Japan) while neuronal activity was monitored.

### Data analysis

#### Behavioral data

To evaluate the degree of interest or attention for each face stimulus, we measured eye-scan patterns for the first face stimuli (S1) that the animals freely scanned. The gaze was quantified separately for three areas: ‘Eyes’, ‘Face without eyes’, and ‘Outside of the face’ (Fig. 2B). Temporal changes in relative gaze frequency between the three areas were computed for every 50 ms window, in shifting 50 ms bins. To elucidate the effects of animacy and reward information on gaze behaviors, we measured the amount of time that the animals spent looking at the eye region of S1. Then, for each session, we computed an ‘animacy index’ using the following formula: animacy index = (Monkey − Cartoon) / (Monkey + Cartoon), where Monkey and Cartoon are the mean amount of time spent looking at the eye region of the monkey and cartoon stimuli, respectively. We also computed a ‘reward index’: reward index = (Large − Small) / (Large + Small), where Large and Small are the mean amount of time spent looking at the eye region of the face stimuli associated with a large and small reward, respectively. A positive animacy index indicated a longer gaze duration for the eye region of the monkey versus cartoon stimuli, and a positive reward index indicated a longer gaze duration for the eye region of the large-reward versus small-reward associated face stimuli.

**Figure 2.**
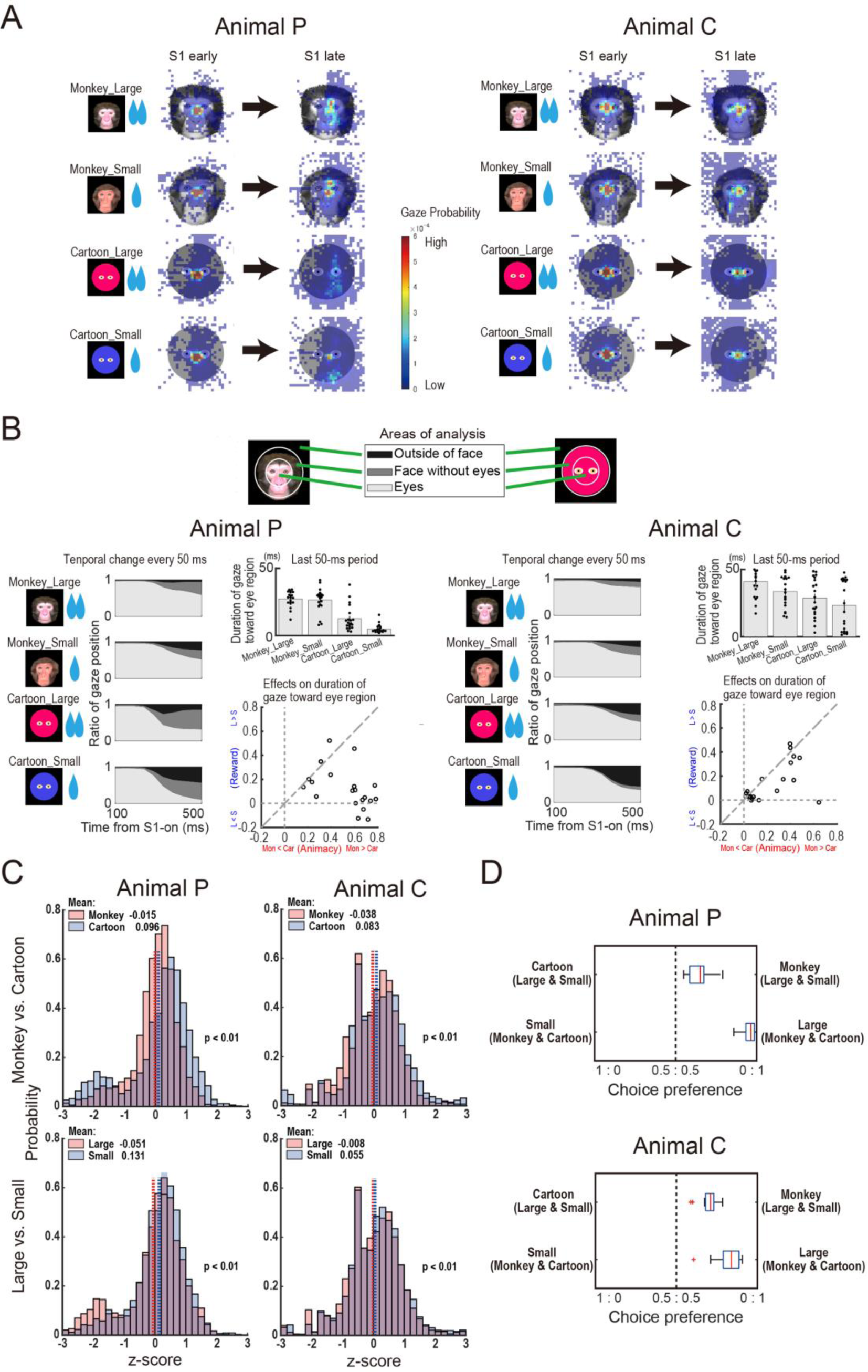
**A.** Persistent focus of attention toward eye region of the monkey face stimuli. Heat maps of each animal’s gaze during the early and late half of the 500-ms S1 presentation period are shown for the four types of visual stimuli: Monkey_Large, Monkey_Small, Cartoon_Large, and Cartoon_Small. **B**. Left: temporal change in gaze position during the S1 presentation period. We analyzed gaze position for three areas: ‘Eyes’, ‘Face without eyes’, and ‘Outside of face’. The ratio of time spent in each area was calculated every 50 ms. Upper right: time spent looking around the eyes during the last 50-ms of S1. Lower right: animacy index (Monkey – Cartoon) / (Monkey + Cartoon) on the x-axis, and reward index (Large– Small) / (Large + Small) on the y-axis, for duration of gaze toward eye region of the S1 (see Methods). **C**. Saccade reaction time distributions. Upper: saccades after the presentation of monkey and cartoon faces. Lower: saccades after the presentation of faces associated with a large and small reward. Reaction times are z-normalized according to the mean and standard deviation of the reaction times for leftward and rightward saccades (see Methods). **D**. Preferences for monkey and large-reward-associated faces. Preferences were determined via a two-alternative forced choice procedure as part of the stimulus-assessment task (Supplementary Figure 2-1). The chosen face in the assessment task was used as the stimulus in the following visually guided saccade procedure. Values on the x-axis denote the preference of each stimulus such that ‘1:0’ denotes a complete preference for the stimulus and ‘0.5:0.5’ denotes equal preference for the stimuli. Box plots indicate median and 25^th^–75^th^ percentiles.

We also analyzed saccade reaction times (SRTs), defined as the interval between the S2 offset and the time at which the velocity of the eye movements exceeded 100 degrees/s. To compare data across the different recording sessions, we converted the raw SRTs to z-scores according to the mean and standard deviation of the SRTs for each direction.

#### Neuronal data

We defined the baseline for each neuron as the mean firing rate during the 500-ms period immediately before the Fix1 onset (Base). Excitatory or inhibitory responses in a given time window were defined as those that were significantly larger or smaller than the signal in the baseline period (Wilcoxon signed rank test, *p* < 0.05).

We constructed peristimulus time histograms (PSTHs) with 1-ms non-overlapping bins and convolved the data with a Gaussian kernel with a standard deviation of 30 ms. For population activity, each neuronal activity was depicted as a z-score that was computed using the mean and standard deviation of the activity during the baseline period.

To elucidate the effects of animacy and reward information on the responses of each amygdala neuron, we computed the strength of the neuronal response to S1 (200 to 700 ms after the onset of S1) and to S2 (100 to 400 ms after the onset of S2) (yellow areas in Fig. 4). For each neuron, we computed an ‘animacy index’ using the following formula: animacy index = (Monkey − Cartoon) / (Monkey + Cartoon), where Monkey and Cartoon are the mean neuronal responses to the monkey and cartoon face, respectively. We also computed a ‘reward index’: reward index = (Large – Small) / (Large + Small), where Large and Small are the mean neuronal responses to the face stimuli associated with a large and small reward, respectively. A positive animacy index value indicated a stronger neuronal response to a monkey face compared with a cartoon face, and a positive reward index value indicated a stronger response to a large-reward associated face compared with a small-reward associated face. We conducted a two-way analysis of variance (ANOVA) for animacy and reward amount. All analyses were performed offline using custom-made Matlab (Mathworks, MA, USA) programs.

**Figure 3.**
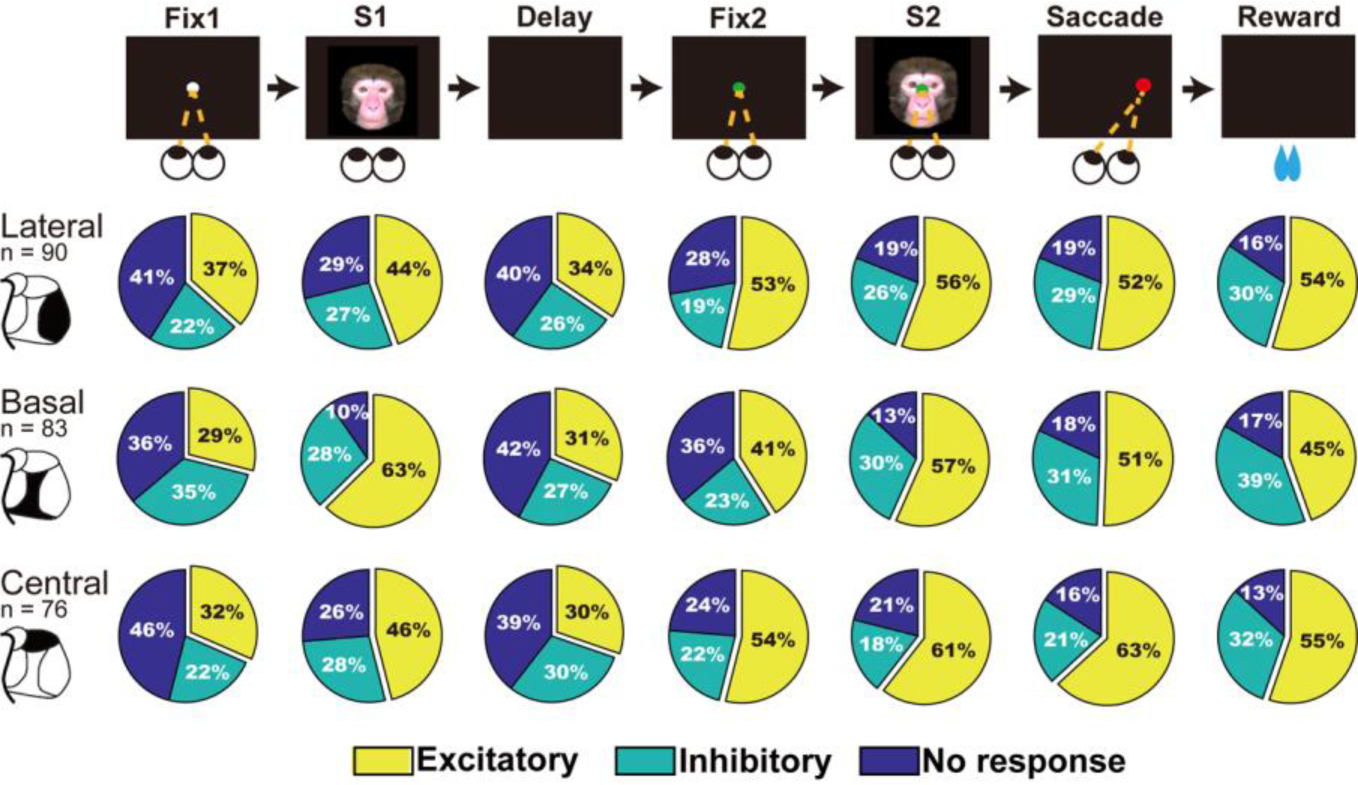
The ratio of neurons with significantly larger excitatory or inhibitory responses during each task period, compared with the baseline activity measured 500 ms before the Fix1.

**Figure 4.**
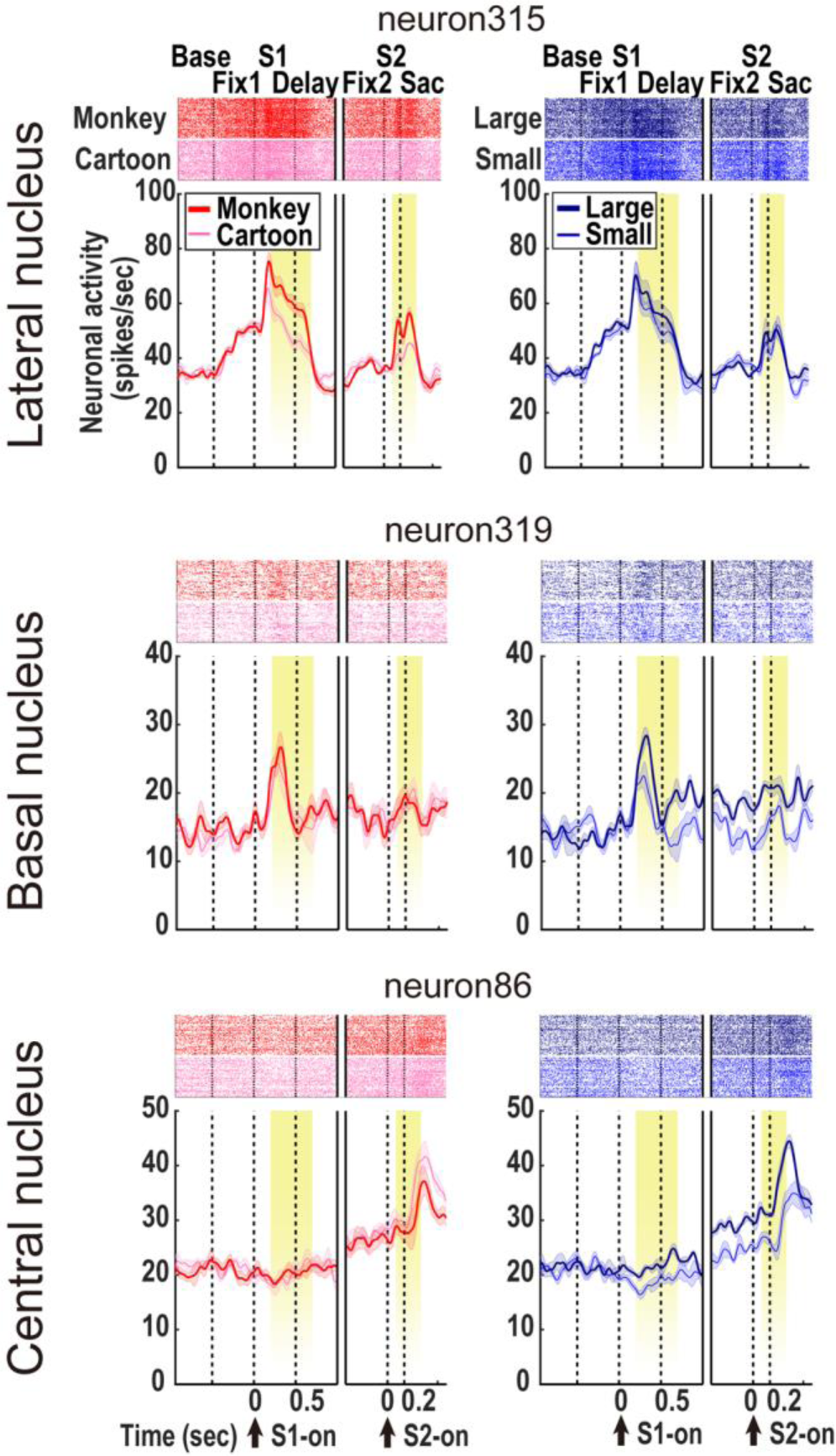
Examples of excitatory single-neuron responses to the S1 and S2 showing animacy or reward information in different amygdala nuclei. We compared the neuronal activity elicited by the monkey versus cartoon face stimuli (animacy effect, left column) or the face stimuli associated with the large versus small reward (reward effect, right column). The yellow area in each diagram indicates the periods considered in the analyses.

To estimate the degree of discrimination for animacy and reward information according to population activity during the task, we performed receiver operating characteristic (ROC) analysis and computed the area under the curve (AUC) for trials involving animacy information (monkey versus cartoon face) or reward information (large versus small rewards) (Fig. 6A). For the animacy information, an AUC value over 0.5 denoted that neurons showed different responses between monkey faces and cartoon faces. For the reward information, an AUC value over 0.5 denoted that neurons exhibited different responses between the large and small reward conditions. To analyze temporal changes in the effects of the animacy and reward information, we calculated the AUC values using sliding windows (200 ms windows with 1 ms steps).

**Figure 5.**
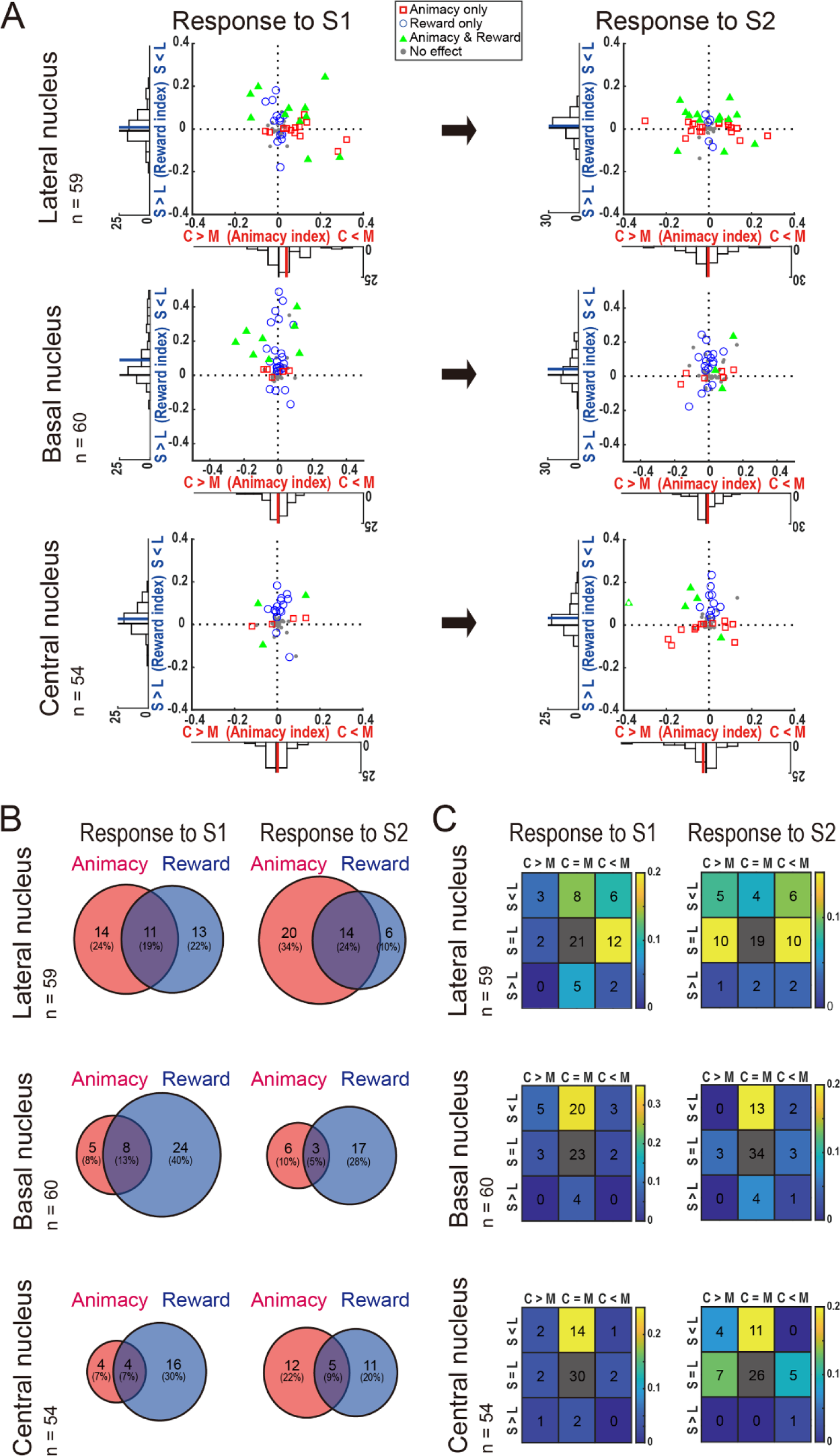
**A.** The animacy index (Monkey – Cartoon) / (Monkey + Cartoon) and reward index (Large– Small) / (Large + Small) for neuronal responses to the S1 (left column) and S2 (right column) (see Methods). Neurons with a significant effect for animacy only, reward size only, or both (p < .05, two-way ANOVA) are shown as red squares, blue circles, and green triangles, respectively. Red and blue lines on the histograms denote the mean animacy index and the mean reward index, respectively. S1, stimulus 1; S2, stimulus 2. **B**. Segregated distribution of neurons that significantly discriminated the degree of animacy and/or reward size (p < .05, two-way ANOVA) in the amygdala subnuclei. The numbers of neurons affected by the animacy factor, reward factor, or both are visualized as Venn diagrams. **C.** the numbers of neurons showing the animacy effect (M > C indicates a significantly stronger response to monkey versus cartoon faces) and reward effect (L > S indicates a significantly stronger response to faces associated with a large versus small reward). The ratios of the numbers of neurons, relative to the neurons modulated by either the animacy or reward factors, are color-scaled.

**Figure 6.**
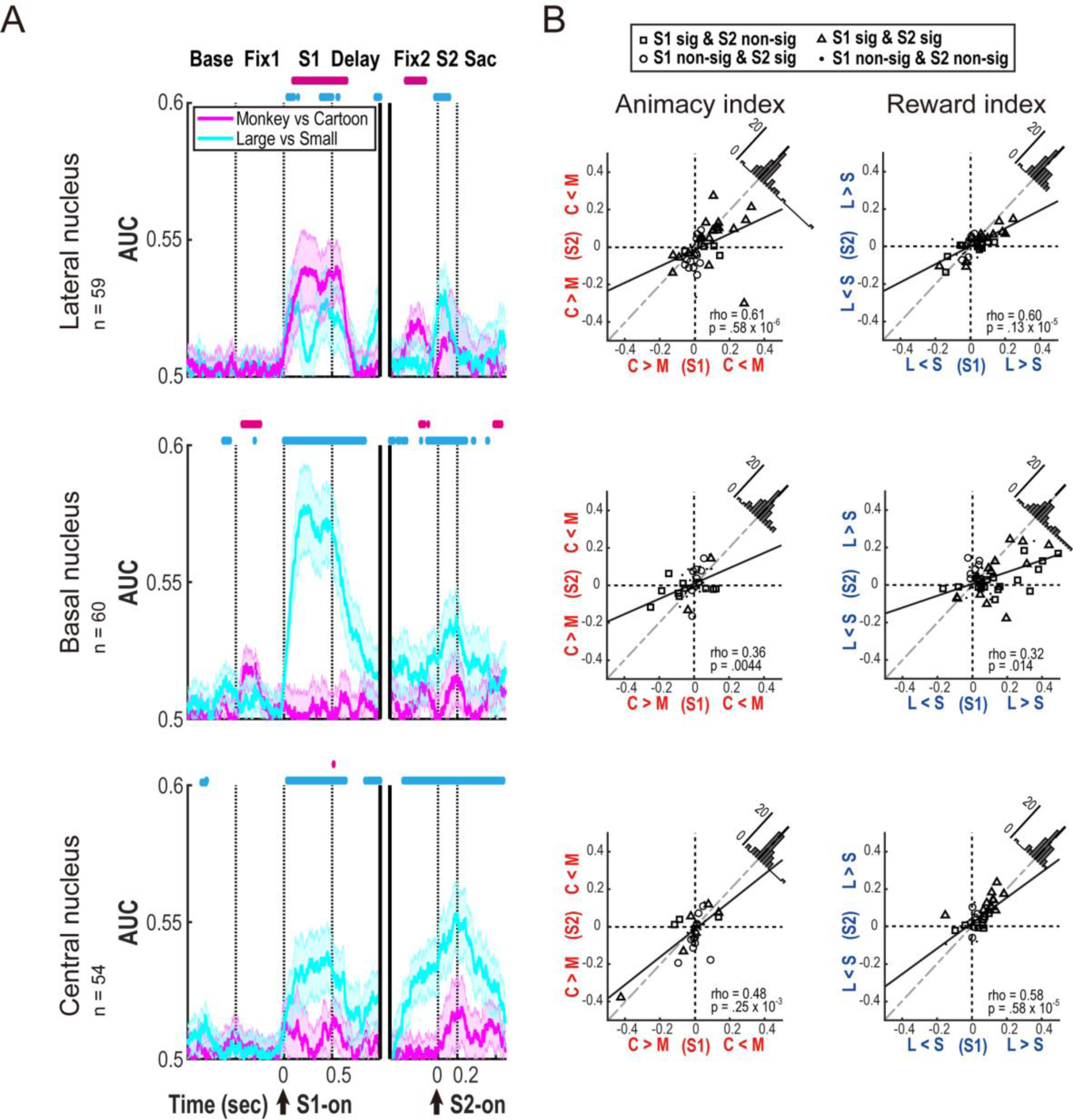
**A.** Time course of animacy- and reward-based signals in neuron populations showing excitatory responses to the S1 and S2 in the lateral, basal, and central nuclei. The AUC (area under the curve) values of the receiver operating characteristic (ROC) analysis enabled us to compare responses to monkey versus cartoon faces (Monkey vs. Cartoon, magenta) or large versus small rewards (Large vs. Small, cyan). A value over 0.5 denotes the degree of discrimination between monkey and cartoon faces or to large and small rewards. The magenta and cyan dots in the upper part of the graphs denote the time points at which the AUC values for animacy and reward, respectively, are significantly different from 0.5. **B**. Consistent animacy (left column, M > C indicates a significantly stronger response to monkey versus cartoon faces) and reward (right column, L > S indicates a significantly stronger response to faces associated with a large versus small reward) information across the S1 and S2 periods. We plotted the animacy and reward indexes of each neuron during the S2 period against those indexes during the S1 period. Neurons with a significant animacy or reward effect in S1 only, S2 only, or both (p < .05, two-way ANOVA) are shown by squares, circles, and triangles, respectively. The inserted histograms show the distributions of the index markers across the diagonal dashed lines.

## Results

### Behavioral data

We recorded single neuronal activity from the amygdala in two animals as they performed a visually-guided saccade task to obtain a liquid reward. In the task, different face stimuli were presented twice (Fig. 1). The visual stimuli had different degrees of animacy (real-monkey or cartoon faces) and different expected reward values (large or small) (Fig. 1A). The stimulus presentation took place immediately after the first fixation period (S1) and just before saccade execution (S2) (Fig. 1B).

The visual stimuli influenced the eye-scan patterns during the presentation of S1, as shown in Figure 2A. Both animals gazed at all of the facial visual stimuli, especially in areas around the eyes, during the first half of the S1 period (Fig. 2A, S1 early). During the later period of S1 (Fig. 2A, S1 late), the gaze was held around the eye area of the monkey-face stimuli or the large reward-associated stimuli. However, the gaze for the cartoon stimuli or small reward-associated stimuli became scattered.

Population data regarding chronological changes in gaze behavior (20 sessions for both animals) support this trend (Fig. 2B, left). For the cartoon compared with the monkey face stimuli, the animals shifted their gaze away from the area around the eyes more often. Especially during the last 50-msec period of S1, the gaze duration for the area around the eyes of the monkey face stimuli was significantly longer than that for the cartoon faces in both animals (Fig. 2B, upper right panel, two-way ANOVA for differences between monkey versus cartoon faces and large versus small rewards; for animal P, F(1, 77) = 121, *p* = .19 × 10^-18^, for animal C, F(1, 77) = 12.8, *p* = .61 × 10^-3^). For small-versus large-reward related stimuli, the gaze was more likely to shift away from the area around the eyes. During the last 50-msec period of S1, the gaze duration for the area around the eyes for the large-reward related stimuli was significantly longer (for animal P, F(1, 77) = 6.83, *p* = .11 × 10^-1^, for animal C, F(1, 77) = 4.00, *p* = .49 × 10^-1^) than that for the small-reward related stimuli (Fig. 2B, upper right panel). These gaze behaviors indicate that the real-monkey and large-reward-associated facial stimuli drew stronger and more persistent attention.

We also compared the strength of the effects of animacy and reward on looking behaviors around eye region of the face stimuli. For each session, we computed an ‘animacy index’, i.e., the relative effect of monkey vs. cartoon faces, plotted on the x-axis, and a ‘reward index’, i.e., the relative effect of large vs. small rewards, plotted on the y-axis (Figure 2B, lower right panel). Comparison of these indices revealed that the animacy effect was significantly stronger than the reward effect in both monkeys (Wilcoxon signed-rank test; for animal P, z = 3.62, *p* = .29 × 10^-5^; for animal C, z = 2.69, *p* = .72 × 10^-2^). This was also evident according to the distribution of the data under the diagonal line.

SRTs to the target were also significantly influenced by both animacy and the expected reward value of the stimuli. SRTs were significantly shorter after presentation of the real-monkey faces versus cartoon faces (two-sample *t*-test; for animal P, *t*(3462) = 3.54, *p* = .40 × 10^-5^; for animal C, *t*(5531) = 5.03, *p* = .51 × 10^-8^; Fig. 2C, upper row). Moreover, SRTs were shorter after the presentation of stimuli associated with large versus small rewards (two-sample *t*-test; for animal P, *t*(3462) = 5.82, *p* = .66 × 10^-10^; for animal C, *t*(5531) = 2.59, *p* = .95 × 10^-4^; Fig. 2C, lower row). These results suggest that both animals paid more persistent and stronger attention to the monkey faces and large-reward-associated faces than the cartoon faces and small-reward-associated faces.

This biased attention or preference was also supported by the task performance in the stimulus-assessment forced-choice task (Supplementary Fig. 2-1). Both animals chose the monkey faces significantly more frequently than the cartoon faces (chi-square test; for animal P, χ^2^ (1, N = 872) = 64.96, *p* = .76 × 10^-17^; for animal C, χ^2^ (1, N = 1277) = 170.78, *p* = .50 × 10^-40^; Fig. 2D,). They also chose the large-reward associated faces more often than the small-reward associated faces (chi-square test; for animal P, χ^2^ (1, N = 849) = 689.31, *p* = .63 × 10^-153^; for animal C, χ^2^ (1, N = 1180) = 429.61 *p* = .20 × 10^-96^; Fig. 2D).

These results indicate that the animals more strongly attended to (or preferred) the real-monkey and large-reward-associated faces compared with the cartoon and small-reward-associated faces. Furthermore, the effect of face type was stronger than the effect of expected reward amount.

### Neuronal activity

#### General

For our neuronal survey, the electrode was directed laterally at an angle of 10 degrees from the dorsal-to-ventral part of the amygdala (Methods, Fig. 9A). After unbiased collection of single neuronal activity data, we identified the affiliation of each task-related neuron (i.e., those with a significantly different response with respect to the baseline period, Methods) to the amygdala sub-nuclei by overlaying penetration record maps on MRI images. We analyzed 90, 83, and 76 single neurons from the lateral, basal, and central nuclei of the amygdala in two animals, respectively. Of these, 64 and 73 in the lateral, 75 and 72 in the basal, and 56 and 60 in the central nuclei showed significant excitatory or inhibitory responses (Methods) to the S1s and S2s, respectively. Evaluating changes in the proportion of responsive neurons over time (Fig. 3) indicated that 1) neurons showing excitatory responses to the S1s or S2s were larger in number than inhibitory-responsive neurons in all sub-nuclei, and 2) the proportion of excitatory responsive neurons increased during the task periods after the second fixation period (Fix2 and later in Fig. 3). We found excitatory responses to S1 or S2 in 59, 60, and 54 neurons in the lateral, basal, and central nuclei, respectively. We found 36, 30, and 24 neurons with inhibitory responses in the lateral, basal, and central nuclei. However, animacy or reward size did not appear to modulate activity in the inhibitory responsive neurons except for reward modulation in the central nucleus (Supplementary Fig. 3-1).

**Figure 7.**
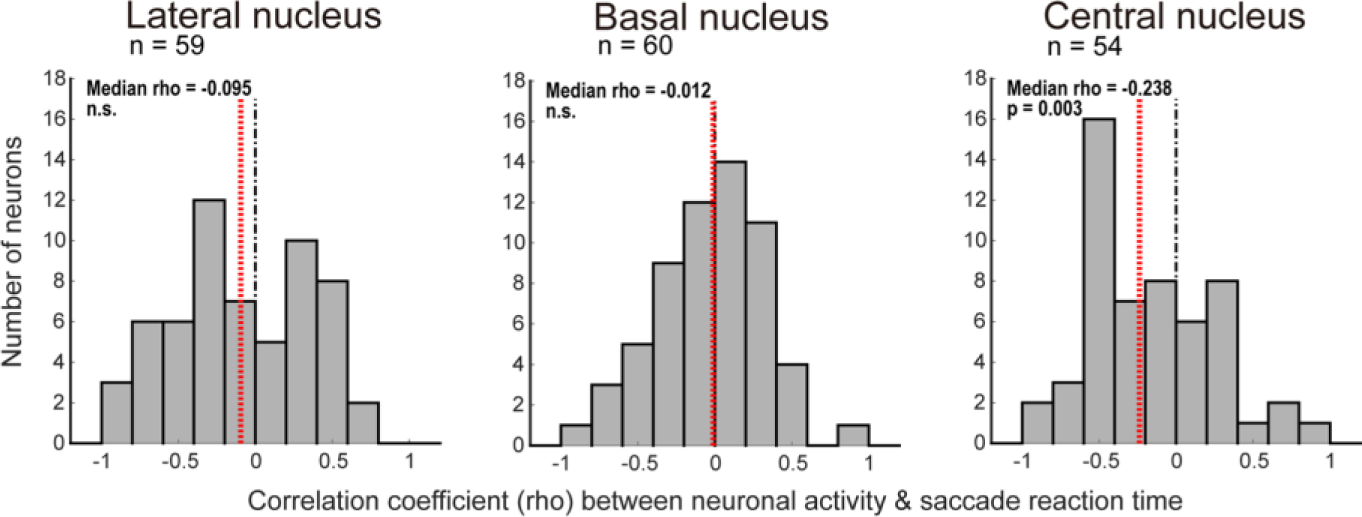
Distribution of Spearman’s rank correlation coefficients between the mean neuronal responses to each stimulus during the S2 and the mean saccade reaction times (SRTs) after each stimulus. Red and black lines on the histograms denote the mean SRTs and 0 of the correlation coefficients, respectively.

**Figure 8.**
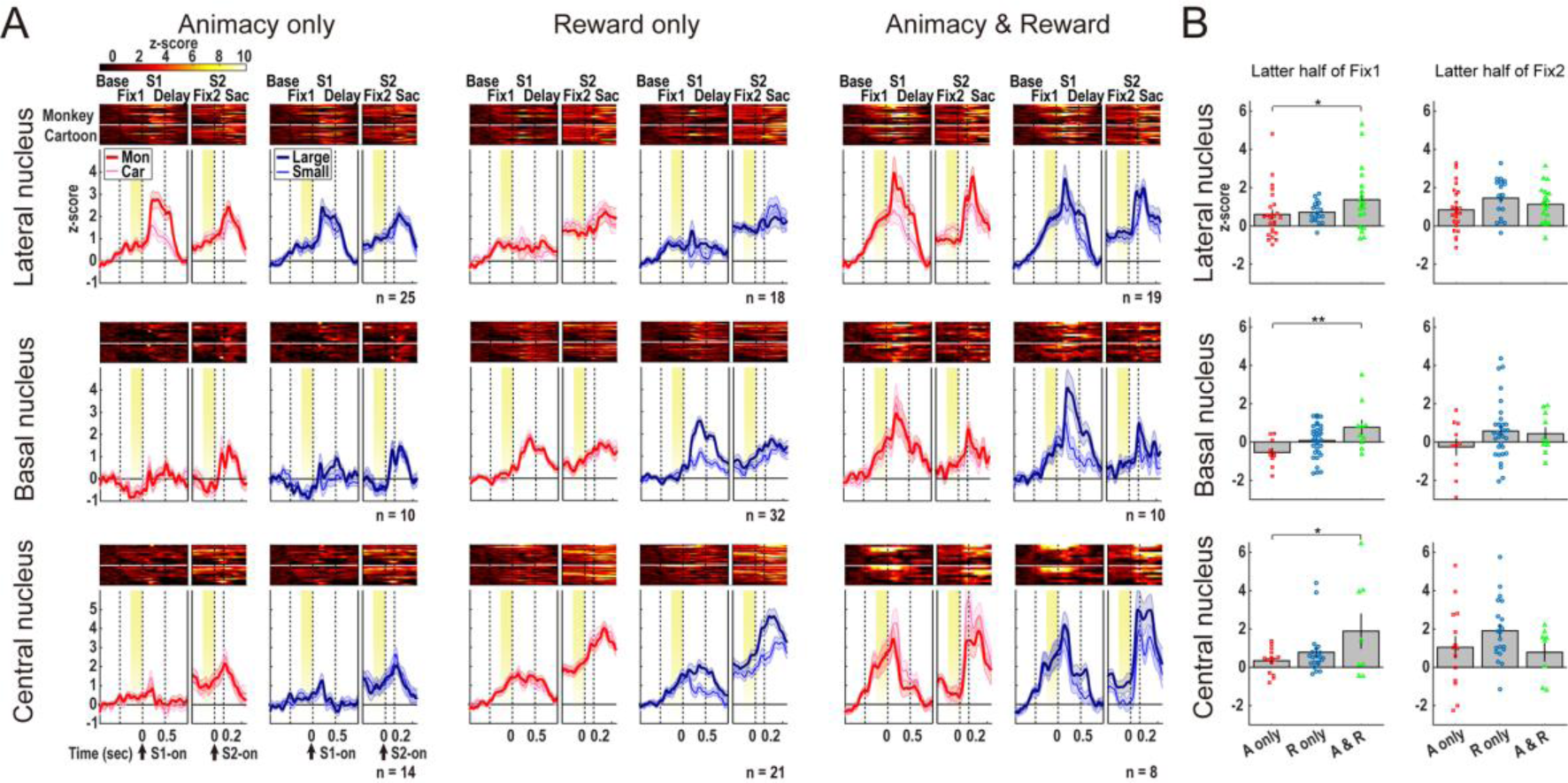
A. The population neurons that discriminated the degree of animacy only, reward size only, and both (p < .05, two-way ANOVA) showed characteristic temporal dynamics. The activity of each neuron is presented as a row of pixels above the histograms. The yellow area in each diagram indicates the periods used in the analyses in (B). B. Mean normalized neuronal activity in the latter half of the Fix1 and Fix 2 periods. Asterisks denote significant differences (*: p < .05, **: p < .01, one-way ANOVA with post hoc Tukey test). Error bars, 1 standard error. A, animacy; R, reward.

**Figure 9.**
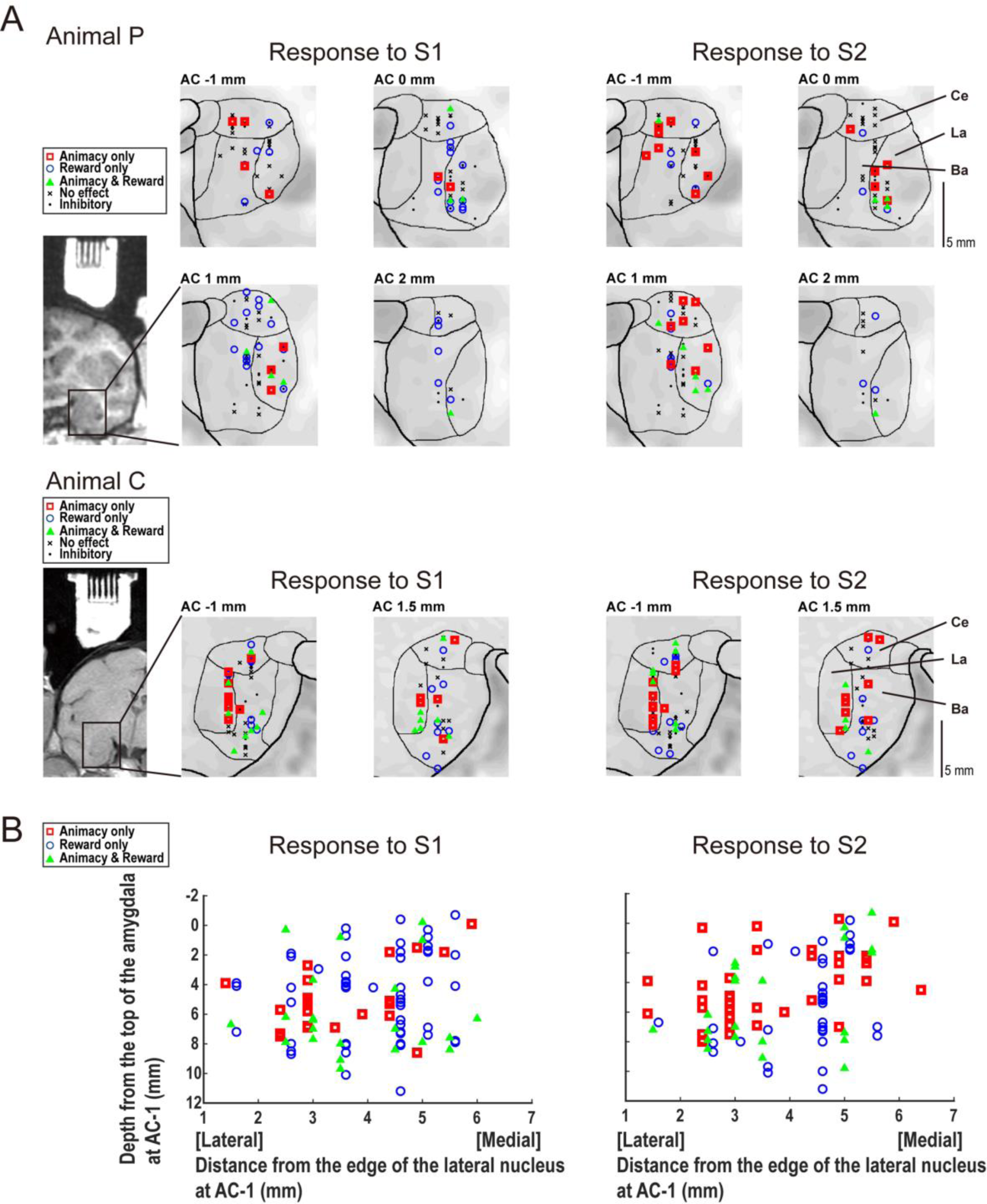
**A.** Estimated locations of the recorded neurons in two monkeys, plotted onto serial coronal sections of MRI images. The data are displayed in anteroposterior order, from 1 mm posterior to 2 mm anterior for Animal P, and from 1 mm posterior to 1.5 mm anterior for Animal C, to the anterior commissure (AC), respectively. Excitatory responsive neurons with a significant effect of animacy only, reward only, or both (p < .05, two-way ANOVA) and non-significant effects of both animacy and reward are shown by red squares, blue circles, green triangles, and black crosses, respectively. Inhibitory responsive neurons are shown by small dots. La, lateral nucleus; Ba, basal nucleus; Ce, central nucleus. B. Mediolateral and dorsoventral distribution of neurons with different effects. The data from all the coronal sections of both monkeys are included. The x-axis indicates the distance from the edge of the lateral nucleus at the AC -1 section. The y-axis indicates the depth from the top of the amygdala at the AC -1 section.

#### Neurons in distinct amygdala sub-nuclei encode animacy and/or reward value according to task context

First, we investigated whether information about animacy and reward size was encoded separately by neurons in the distinct amygdala sub-nuclei. Figure 4 shows representative examples of the activity of a single neuron in each sub-nucleus. Our comparison of the responses to the monkey- and cartoon-faces (Fig. 4, left column) revealed stronger responses to the monkey faces following S1 and S2 in the lateral nucleus neuron (Fig. 4, upper left panel). We found no differences in the responses to S1 and S2 in the basal nucleus neuron (Fig. 4 middle left panel). The central nucleus neuron exhibited stronger responses to the cartoon faces versus the monkey faces following S2 (Fig. 4, lower left panel). By contrast, a comparison between the responses to the large- and small-reward stimuli (Fig. 4, right column) revealed stronger responses for large reward-versus small reward-associated faces during the S1 and S2 periods in the basal and central nuclei neurons (Fig. 4, middle and lower right panels), although there was no difference in the lateral nucleus neuron (Fig. 4, upper right panel). These results indicate that distinct amygdala sub-nuclei differentially encode animacy- and reward-related information during specific task epochs. We also identified neurons with other response types (Supplementary Figure 4-1).

To further investigate how individual neurons process animacy and reward information, we computed the ‘animacy index’, i.e., the relative encoding strength of monkey- vs. cartoon-faces, and the ‘reward index’, i.e., the relative encoding strength of large- vs. small-rewards (Methods). Figure 5A, shows the animacy index on the x-axis and reward indices on the y-axis for each neuron.

For the response to S1 in the lateral nucleus (Fig. 5A, upper left panel), many data points were located in the right half of the graph (also see the histogram at the bottom), indicating stronger responses to monkey-versus cartoon-face stimuli. The mean animacy index was 0.0367 ± 0.0115 (mean ± SEM), which is significantly higher than 0 (Wilcoxon signed-rank test, z = 3.00, *p* = .27 × 10^-2^ ). The data plots were distributed equally around 0 along the y-axis (0.0054 ± 0.0147, z = 1.20, *p* = .23). This indicates either positive or negative reward coding by individual neurons during S1. For S2, the data points were distributed widely along the x-axis, with an equal number of points in the positive and negative directions. This indicates positive or negative coding of animacy by individual neurons. The distribution on the y-axis was narrow, indicating a small degree of reward coding. We also found that 11 and 14 neurons during the S1 and S2 periods, respectively, encoded significant effects of both animacy and the reward effect (p < .05, two-way ANOVA), as shown by green triangles. Thus, a group of lateral-nucleus neurons encoded multiple different types of information.

For the response to S1 in the basal nucleus (Figure 5A, middle left), most of the data points were in the upper half of the graph, while the distribution along the x-axis was small, indicating a stronger reward signal. A similar trend was observed for the response to S2 (Figure 5A, middle right), although the strength of the reward signal was smaller than that for S1. The mean reward indexes for the responses to S1 and S2 were 0.0894 ± 0.0179 and 0.0344 ± 0.0111 (mean ± SEM), respectively, which is significantly higher than 0 (z = 4.84, *p* = .31 × 10^-7^ and z = 2.59, *p* = .96 × 10^-2^, respectively). Conversely, the mean animacy indexes for the responses to S1 and S2 were not significantly different from 0. Some basal nucleus neurons also showed significant modulation by both animacy and reward during the S1 period.

Neurons in the central nucleus showed dominant reward coding during the S1 and S2 periods (Figure 5A, lower panel), which was similar to our data for the basal nucleus. The mean reward indexes for the responses to S1 and S2 in the central nucleus were 0.02242 ± 0.0087 and 0.0335 ± 0.0088 (mean ± SEM), respectively, which is significantly higher than 0 (z = 3.05, *p* = .23 × 10^-2^ and z = 3.39, *p* = .70 × 10^-3^, respectively). The mean animacy indexes for the responses to S1 and S2 were not significantly different from 0.

These results indicate that the effects of animacy information were dominant over those of reward information in the lateral nucleus, whereas the effects of reward information were dominant over those of animacy information in the basal and central nuclei.

Next, we classified the neurons that were responsive to the S1 and/or S2 in each sub-nucleus, based on a two-way analysis of variance with animacy and reward as variables. Figure 5B shows the number of neurons that were significantly affected by the animacy (red circles) and/or reward factors (blue circles). In terms of the response to the S1 and S2, we found more neurons that were affected *solely* by the animacy or reward factor compared with those affected by *both* factors in almost all cases for all sub-nuclei (chi-square test; for the response to the S1 in the lateral nucleus, χ^2^ (1, N = 38) = 6.74, *p* = .94 × 10^-2^; for the response to the S1 in the basal nucleus, χ^2^ (1, N = 37) = 11.92, *p* = .55 × 10^-5^; for the response to the S2 in the basal nucleus, χ^2^ (1, N = 26) = 15.28, *p* = .87 × 10^-6^; for the response to the S1 in the central nucleus, χ^2^ (1, N = 24) = 10.67, *p* = .11 × 10^-2^; for the response to the S2 in the central nucleus, χ^2^ (1, N = 28) = 11.57, *p* = .67 × 10^-5^) except for the response to the S2 in the lateral nucleus ( χ^2^ (1, N =40) = 3.60, *p* = .58 × 10^-1^).

In Figure 5C, we further classified neurons depending on preference for face-type or reward size. In the lateral nucleus, we found more neurons with stronger responses to monkey versus cartoon faces compared with the number of neurons with stronger responses to cartoon faces versus monkey faces during S1 (chi-square test, χ^2^ (1, N = 25) = 9, *p* = .27 × 10^-2^). This trend, however, changed during S2: the numbers of neurons that preferred monkey faces and cartoon faces were nearly equal.

In the basal and central nuclei, during both the S1 and S2 periods, we found more neurons with stronger responses to the large-reward versus small-reward associated faces (for the response to the S1 in the basal nucleus, χ^2^ (1, N = 32) = 18.00, *p* = .22 × 10^-6^; for the response to the S2 in the basal nucleus, χ^2^ (1, N = 20) = 5.00, *p* = .25 × 10^-1^; for the response to the S1 in the central nucleus, χ^2^ (1, N = 20) = 9.80, *p* = .17 × 10^-2^; for the response to the S2 in the central nucleus, χ^2^ (1, N = 16) = 12.25, *p* = .47 × 10^-5^).

#### Animacy and reward value coding were consistent but varied in strength across different task epochs

The variations in the impact of the animacy and reward information during S1 and S2 (Fig. 5) indicated the effect of their temporal positions during each trial. For further quantitative analyses, for each neuron, we computed AUC values to compare the activity within sliding 200-ms windows regarding animacy or reward (Methods), and averaged the AUC values separately for each amygdala nucleus (Fig. 6A).

In the lateral nucleus, the animacy signal (magenta curves) increased and remained at a high level during presentation of the S1, confirming that the response discriminated monkey faces from cartoon faces. The animacy signal then decreased during and after the delay period, including the S2 period. Although the strength of the animacy signal during S1 and S2 was different when the neurons were analyzed as a population (Fig. 6A, upper panel), the effect of the animacy information remained consistent in the majority of the neurons, as revealed by a significant positive correlation between the animacy index for S1 (x-axis) and S2 (y-axis) (Fig. 6B upper left panel, Spearman’s rank correlation coefficient, rho = .61, p = .58 × 10^-6^). The orthogonal distribution was biased toward the area under the diagonal line (Wilcoxon signed-rank test, z = 2.70, *p* = .69 × 10^-2^), indicating a stronger animacy signal for S1 versus S2. Note, however, that some neurons showed changes in the animacy signal between S1 and S2. For example, in Fig. 6B, upper left panel, the signal was weak during S1 but large during S2 (circles, n = 13) or opposite (squares, n = 4). This explains the bilateral animacy preference observed during S2 (Fig. 5C, Lateral, S2).

In the lateral nucleus, the reward signal (cyan curves in Fig. 6A, upper panel) increased and stayed at a moderate level during S1. The reward signal then reduced and increased again during the S2 period. At the single neuron level, the reward information for each neuron was kept constant, as revealed by a significant positive correlation between the reward index for S1 (x-axis) and S2 (y-axis) (Fig. 6B upper right panel, rho = .60, *p* = .13 × 10^-5^).

In the basal nucleus, the reward information was prominently high during S1 and continuously moderate during the subsequent task periods (cyan curves in Fig. 6A, middle panel). The positive correlation between the reward index for S1 and S2 (rho = .32, *p* = .014, Fig, 6B, middle right panel) indicates that the reward information was consistent across S1 and S2. Most data points corresponding to the reward indices were under the diagonal line, indicating that the reward signals were stronger during the S1 versus S2 period in many basal neurons (z = 2.94, *p* = .33 × 10^-2^). In the basal nucleus, the animacy signal did not show significant modulations throughout the trial (Fig. 6A, middle panel).

In the central nucleus, the reward information was continuously high during S1, the delay, and S2, and continued to be at a high level even after the S2 disappeared (cyan curves in Fig. 6A, lower panel). The reward indices for S1 and S2 retained a positive correlation (Fig. 6B, lower right), and there was no bias toward either S1 or S2. The animacy signal did not show signs of significant modulation throughout the trial (Fig. 6A, lower panel).

Although there were few neurons that showed a significant animacy effect in the basal and central nuclei, we found a positive correlation between the animacy signal in S1 and S2 (basal: rho = .36, *p* = .44 × 10^-2^, and central: rho = .58, *p* = .58 × 10^-5^, l: Fig. 6B, middle and lower left panels), indicating consistent coding across task epochs as a neuronal population.

The results shown in Figures 5A and 6B suggest that the animacy information was primarily processed in the lateral nucleus in the early (sensory encoding) phase of the task. Conversely, the reward signal was continuously discriminable throughout trials in all sub-nuclei. However, we found a difference in the temporal modulation pattern: in the basal nucleus, reward information was prominent during the early phase, while in the central nucleus, no difference between task periods was observed.

In the present task, the S2 was presented just before saccade execution. To investigate whether the activity during S2 contributed to saccade execution, we calculated Spearman’s rank correlation coefficients between the mean activity during S2 and the mean SRTs for each neuron. The median correlation coefficients (*ρ*) of the neuronal populations in each sub-nucleus were -0.095 (Lateral), -0.012 (Basal), and -0.238 (Central). The correlation coefficients were significantly biased toward negative values in the central nucleus only (Wilcoxon signed-rank test; z = -2.97, *p* = .30 × 10^-2^; Fig.7). These results suggest that, in the central nucleus rather than the lateral and basal nuclei, stronger activity during S2 was associated with quicker saccades.

#### Amygdala neurons showing both animacy and reward effects represented specific activity patterns during the fixation periods

At this point, our analyses had indicated that animacy and reward information are represented to different degrees with dynamic changes in distinct amygdala nuclei. We further found that neurons with these different signal coding types, i.e., animacy only, reward only, and both animacy and reward, had specific temporal activity patterns.

A noticeable difference in neuronal activity among neurons with distinct signal coding types (Fig. 8A) appeared during the Fix1 period. Neurons that exhibited the animacy effect only (Fig. 8A, left two columns) exhibited slightly increased (lateral and central nuclei) or decreased (basal) activity during Fix1 compared with the baseline. Neurons sensitive to the reward effect only (Fig. 8A, middle two columns) exhibited a slight increase in activity during Fix1 in all nuclei. In contrast, neurons sensitive to both animacy and reward information (Fig. 8A, right two columns) exhibited a large increase in activity throughout the Fix1, peaking during the S1 period. Notably, this pattern was observed across the amygdala subnuclei. The mean neuronal activity during the latter half of Fix1 was the highest (one-way ANOVA; for the lateral nucleus, F(2, 59) = 3.39, *p* = .40 × 10^-1^; for the basal nucleus, F(2, 49) = 5.66, *p* = .62 × 10^-2^; for the central nucleus, F(2, 40) = 3.31, *p* = .47 ×10^-1^) for neurons with both animacy and reward effects (Fig, 8B, left panels of each nucleus). Given that the Fix1 period was *before* the presentation of the visual stimuli, this activity appears to have been associated with the general preparation process. The number of individual neurons exhibiting increased activity during Fix1 also reflected the superiority of neurons sensitive to both animacy and reward information. A Z-score over 2 denotes a significant (*p* < .05) increase in neuronal activity from the baseline. The proportion of neurons showing a Z-score over 2 was 12.0% (3/25), 0% (0/18), and 26.3% (5/19) in the lateral nucleus, 0% (0/10), 0% (0/32), and 20.0% (2/10) in the basal nucleus, and 0% (0/14), 9.5% (2/21), and 37.5% (3/8) in the central nucleus for the animacy effect only, reward effect only, and both the animacy and reward effects, respectively.

The animals also fixated on the central fixation point during the Fix2 period. The activity during Fix2 was high in the reward only-type neurons (neurons showing z-scores over 2, 16.0% (4/25)) in the lateral nucleus and animacy only-type neurons (28.6% (4/14)) and reward only-type neurons (42.9% (9/21)) in the central nucleus, while it was weak in the animacy only-type neurons (16.0% (4/25)) and the both animacy and reward-type neurons (15.8% (3/19)) in the lateral nucleus, all types of neurons (0% (0/10) for the animacy effect only, 15.6% (5/32) for the reward effect only, and 0% (0/10) for both the animacy and reward effects) in the basal nucleus, and the both animacy and reward-type neurons (12.5% (1/8)) in the central nucleus (Fig, 8B, right panels of each nucleus).

#### Localization of neurons with distinct effects

Figure 9A shows the estimated locations of the analyzed neurons (Fig. 3), plotted on a serial of coronal sections of MRI images. In both animals, neurons with a significant animacy effect were widely observed from the dorsal to ventral part of the lateral nucleus. Many neurons with significant reward effects were observed in the central and basal nuclei. We also found that neurons with both animacy and reward effects tended to be distributed in the ventral part of the lateral or basal nuclei. The mediolateral and dorsoventral distribution of the neurons with animacy, reward, and both effects during S1 confirmed these trends (Fig. 9B). While neurons with the reward effect only (blue circles) were scattered overall, neurons with the animacy effect only were located significantly lateral to those with the reward effect only (Wilcoxon rank sum test; for the responses to S1, z = -2.27, *p* = .23 × 10^-1^; for the responses to S2, z = -2.22, *p* = .26 × 10^-1^). Neurons with both the animacy and reward effects were located significantly ventral to those with the animacy effect only (Wilcoxon rank sum test; z = -2.14, *p* = .32 × 10^-1^).

## Discussion

### Primates perceive information about animacy from conspecifics

The eye gaze patterns and SRTs revealed that the animals perceived the real monkey and cartoon face stimuli differently. Although differences in the properties of visual stimuli could affect gaze behavior, in the present study, all visual stimuli had a uniform mean luminance and size. Behavioral differences could also be related to the saliency of visual stimuli. The sustained attention (Fig. 2A and 2B), shorter SRTs (Fig. 2C), and preference (Fig. 2D) for monkey faces indicates that the real face stimuli were more salient. The difference in behavioral results, however, cannot be explained solely by the saliency of the stimuli. In the present task, for the animal subjects, the type of reward (large or small) was expected to be more relevant than the type of stimuli (monkey or cartoon). However, the stimuli type had a greater effect on the eye gaze pattern (Fig. 2B, lower right panel). Altogether, the simple visual properties or degree of saliency of the stimuli could not explain the different behavioral effects of the stimuli type; instead, the behavioral results appear to have been influenced by the animacy of the face stimuli.

Animate stimuli attract more attention than inanimate stimuli (Lindemann et al., 2011). This is consistent with our behavioral findings that monkey faces were more attractive than cartoon ones. Furthermore, monkeys’ perception of facial reality was reported to be similar to that of humans. Macaque monkeys feel eeriness toward an artificial face that looks too much like a real one (Steckenfinger and Ghazanfar, 2009): this phenomenon is called the uncanny valley and is well-known in humans. This evidence indicates that monkeys perceive animacy from a real face. Thus, our animals might have discriminated the degree of animacy between the monkey and cartoon faces.

### Animacy information is encoded differently in distinct amygdala subnuclei

We found that many neurons in the lateral nucleus of the amygdala discriminated face stimuli in terms of animacy. Neurons in the primate lateral and basal nuclei process facial expressions (Gothard et al., 2007; Kuraoka and Nakamura, 2007; Inagaki et al., 2023), gaze direction (Tazumi et al., 2010; Mosher et al., 2014), and the social hierarchy of faces (Munuera et al., 2018). The lateral nucleus receives projections from the area TE and TEO in the inferotemporal (IT) cortex (Ghashghaei and Barbas, 2002; Stefanacci and Amaral, 2002), the dorsal bank of the superior temporal sulcus (STS) (Stefanacci and Amaral, 2000), and the pulvinar (Jones and Burton, 1976; Day-Brown et al., 2010). Of these, neurons in the primate TE and the STS distinguish animate from inanimate images (Kiani et al., 2007; Kriegeskorte et al., 2008; Ninomiya et al., 2021). In contrast, pulvinar neurons show similar responses to real and cartoon faces (Nguyen et al., 2013). Thus, face animacy information in the lateral nucleus may originate from higher-order visual areas, including the TE and STS.

Although most animacy-sensitive neurons in the lateral nucleus responded more strongly to the monkey face stimuli during S1, a similar number of neurons responded more strongly to the monkey or cartoon faces during S2 (Fig. 5A upper right panel, 5C, upper right panel, and Fig. 6B, upper left panel). This change in response preference indicates that the task context might influence the animacy signal. The encoding of animacy information precedes other information processes, and may be an innate response. However, face identity is irrelevant to the task. Indeed, changes in preference were not common for the reward signals (Fig. 6B, upper right panel), which are continuously relevant throughout a trial. Changes in animacy coding were also observed in the central nucleus. While animacy was not observed during S1, neurons that responded significantly more strongly for either cartoon or monkey faces emerged during S2 (Fig. 4, Supplementary Fig.4-1, Fig. 5A and 5C, lowest row). The face animacy signal in the central nucleus during the latter phase of the task might rely on information from the lateral nucleus, given that the central nucleus receives intra-amygdala projections from the lateral nucleus (Pitkänen and Amaral, 1998).

### Reward information is encoded differently in distinct amygdala sub-nuclei

While some studies reported homogenous distribution of reward-coding neurons across the amygdala sub-nuclei (Sugase-Miyamoto and Richmond, 2005; Paton et al., 2006; Belova et al., 2007; Bermudez and Schultz, 2010; Bermudez et al., 2012; Iwaoki and Nakamura, 2022), others reported that the basolateral nucleus is involved in stimulus-reward associations (Fuchs et al., 2006; Feltenstein and See, 2007). Here, we observed the reward effect in all amygdala nuclei, with varying degrees and temporal dynamics.

The reward signal in the lateral and basal nuclei that corresponded to the preference for large versus small reward-related face stimuli was more prominent during S1 versus S2 (Fig. 5A and Fig. 6A, upper and middle panels). This dominant reward signal during S1 indicated that sensory analyses took place in relation to the expected rewards. However, the lateral and basal nuclei had different response patterns. Neurons in the lateral nucleus neurons showed a phasic reward signal, while those in the basal nucleus were continuous during S1, followed by delay and fixation periods. This indicates partially distinct reward circuits. Indeed, while the lateral nucleus receives input from the temporal cortex, especially TE, TEO, and STS (Stefanacci and Amaral, 2000), the basal nucleus has reciprocal connections with the anterior cingulate and orbitofrontal cortex (Ghashghaei and Barbas, 2002; Stefanacci and Amaral, 2002), and receives projections from the lateral nucleus (Pitkänen and Amaral, 1998).

The reward signal in the central nucleus was characterized by a continuous large-reward preference, which was equally strong throughout the trial until saccade onset (Fig. 5A and Fig. 6A, lower panels). The central nucleus receives strong intra-amygdala projections (Price and Amaral, 1981; Pitkänen and Amaral, 1998), and projects to subcortical areas involved in autonomic responses (Price and Amaral, 1981; Jongen-Rêlo and Amaral, 1998; Freese and Amaral, 2009). Thus, this supports the involvement of the central nucleus in autonomic responses. The central nucleus also projects to subcortical areas involved in reward (Price and Amaral, 1981; Jongen-Rêlo and Amaral, 1998; Fudge and Haber, 2000; Freese and Amaral, 2009). Among these, the output from the central nucleus to the basal ganglia circuit is known to modulate saccadic eye movements (Maeda et al., 2020; Maeda et al., 2023). Indeed, SRTs were negatively correlated with neuronal activity in the central nucleus during the S2 (Fig. 7), and this elevated neuronal activity might facilitate saccadic eye movements. Taken together, these data suggest that the central nucleus integrates information from other brain regions to enable action execution.

### Integrated and separate encoding of animacy and reward information

We also found a group of neurons that encoded *both* face animacy and reward information. The amygdala basolateral nuclei reportedly exhibit shared parallel coding of social and valence (reward or punishment) information at the neuronal population level, such as social hierarchy and reward size (Munuera et al., 2018), or gaze direction and valence (Pryluk et al., 2020). In contrast, individual neurons mostly processed either social or valence information (Putnam and Gothard, 2019). Notably, we further identified unique features of multi-coding neurons across different amygdala nuclei.

First, the neurons with both animacy and reward coding showed characteristic buildup activity during Fix1. Amygdala activity during the pre-stimulus fixation period reportedly encodes a positive state value (Belova et al., 2008). Similar pre-stimulus activity was reported in the basolateral amygdala (Sugase-Miyamoto and Richmond, 2005), which is potentially related to increased arousal or attention expecting rewards. The buildup acitvity was followed by stronger activity in response to the face stimuli (Fig. 8A). Thus, neurons with both effects may process arousal or attention to future reward and sensory information.

Second, neurons that encode both animacy and reward were primarily found in the ventral lateral and basal amygdala (Fig. 9A, B). This supports the integration of information in the lateral and basal nuclei, given that efferent fibers mainly originate in the dorsal region and terminate in the ventral division (Price et al., 1987; Pitkänen and Amaral, 1998). Thus, animacy or reward information travels from the dorsal to ventral amygdala, where integration of both types of information takes place. Shared encoding of animacy and reward information may partly be explained by common features of the stimuli, such as similar saliency. Alternatively, the shared coding of both types of information may result from the integration of both types of information within the amygdala.

### Conclusion

In conclusion, our data indicate that face animacy is predominantly processed in the lateral nucleus, reward information is predominantly processed in the basal nuclei for sensory analyses, and continuous reward coding takes place in the central nucleus for action execution. Some neurons encode both animacy and reward, probably through intra-nuclei integration. Further studies should examine whether each nucleus plays a causal role in the perception of animacy or reward. Assessing information flow between the subnuclei as well as other non-amygdala brain areas could lead to separate or integrated analyses of social and valence information processing.

## Acknowledgments

This work was supported by a JSPS KAKENHI Grant Number JP19K03388, JP21H00312, JP22K03203, and JP23H03842 (to K.K.); and a JSPS KAKENHI Grant Number JP19H03540, JP21H00216, and JP22K19485; AMED-CREST 21gm1510003 (to K.N.). We are grateful to K. Adachi, M. Arisato, H Shimazaki, H. Onoe, and T. Isa for obtaining magnetic resonance images.

The subject monkeys were provided by NBRP “Japanese Monkeys” through the National BioResource Project of the MEXT, Japan.

We thank Sydney Koke, MFA, from Edanz (https://jp.edanz.com/ac) for editing a draft of this manuscript.

## Supplementary Materials

**Supplementary Figure 1-1.**
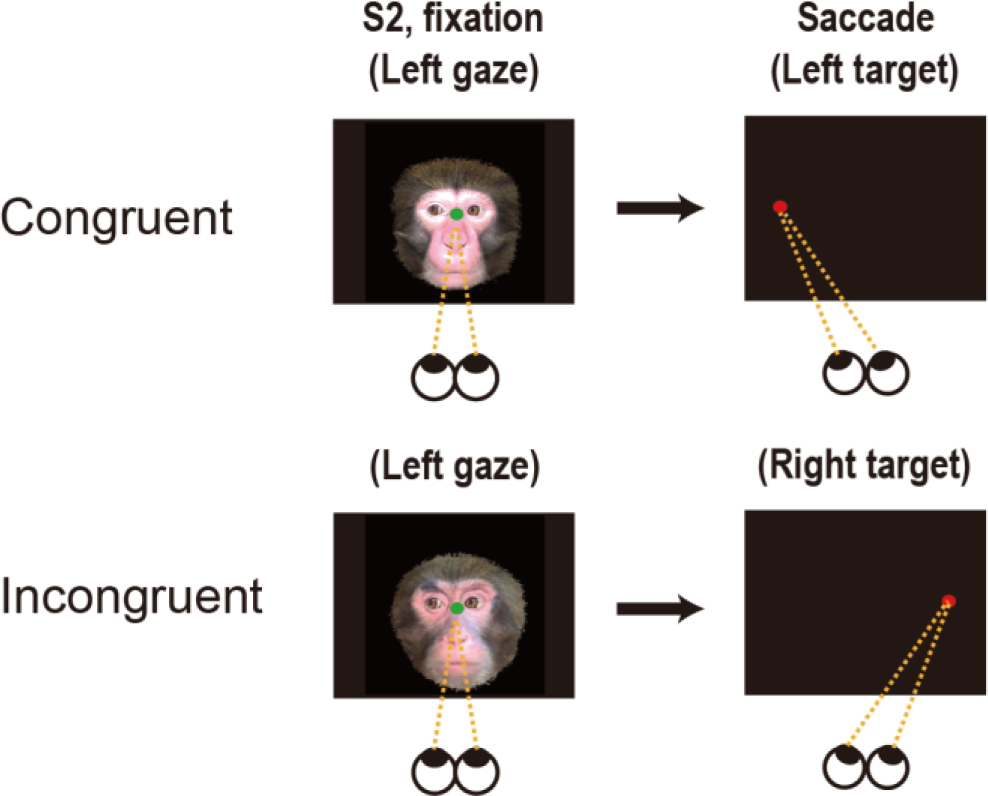
Congruency of the gaze of S2 and the location of the target. The gaze direction of the S2 faces was either left or right. The gaze direction of half of the S2 faces (M1L, M3S, C1L, C3S from Figure 1A) was always congruent with the future target (Congruent). The other half of the S2 faces (M2L, M4S, C2L, C4S) was always incongruent with the future target (Incongruent).

**Supplementary Figure 2-1.**
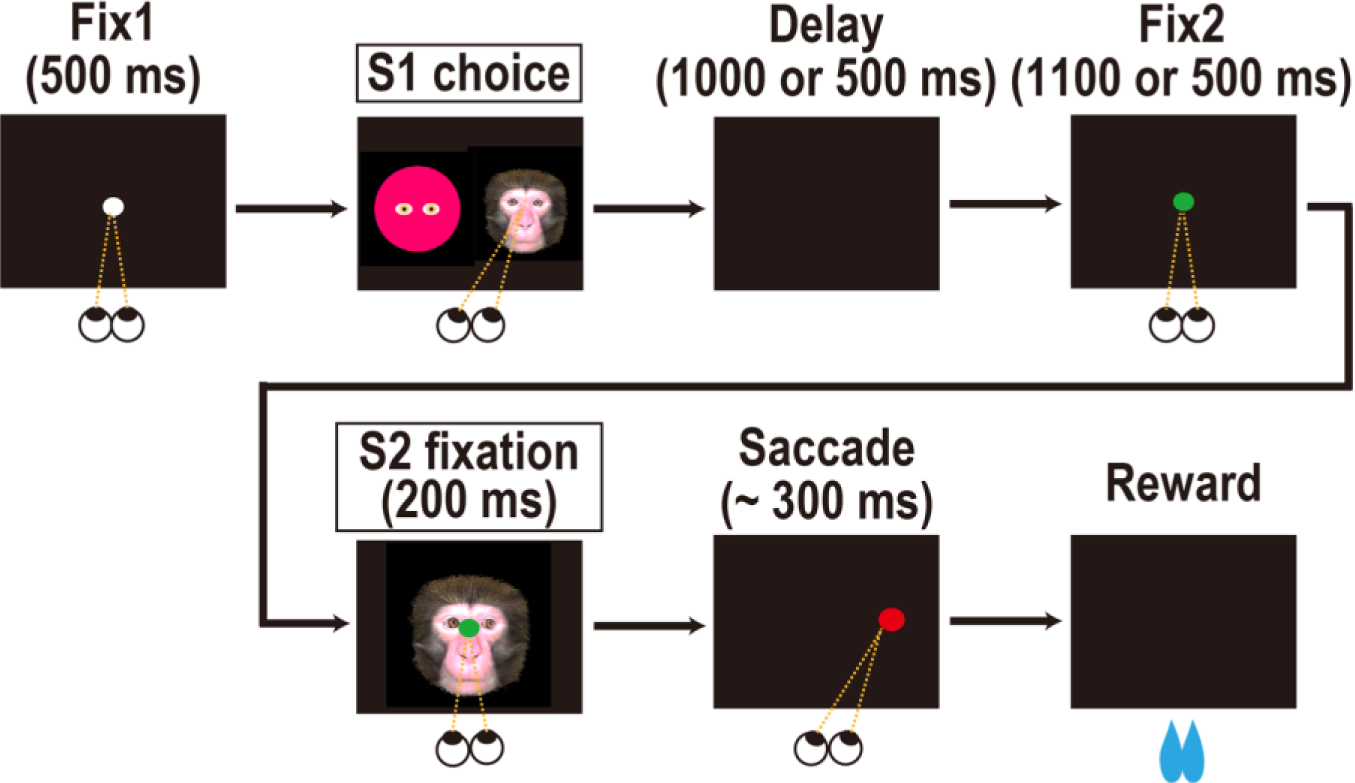
Stimulus-assessment task. The subjects were forced to choose one of two face stimuli simultaneously presented as S1. They chose between a monkey and cartoon face or between large-and small-reward associated faces. The chosen stimulus was presented in the following S2 period.

**Supplementary Figure 3-1.**
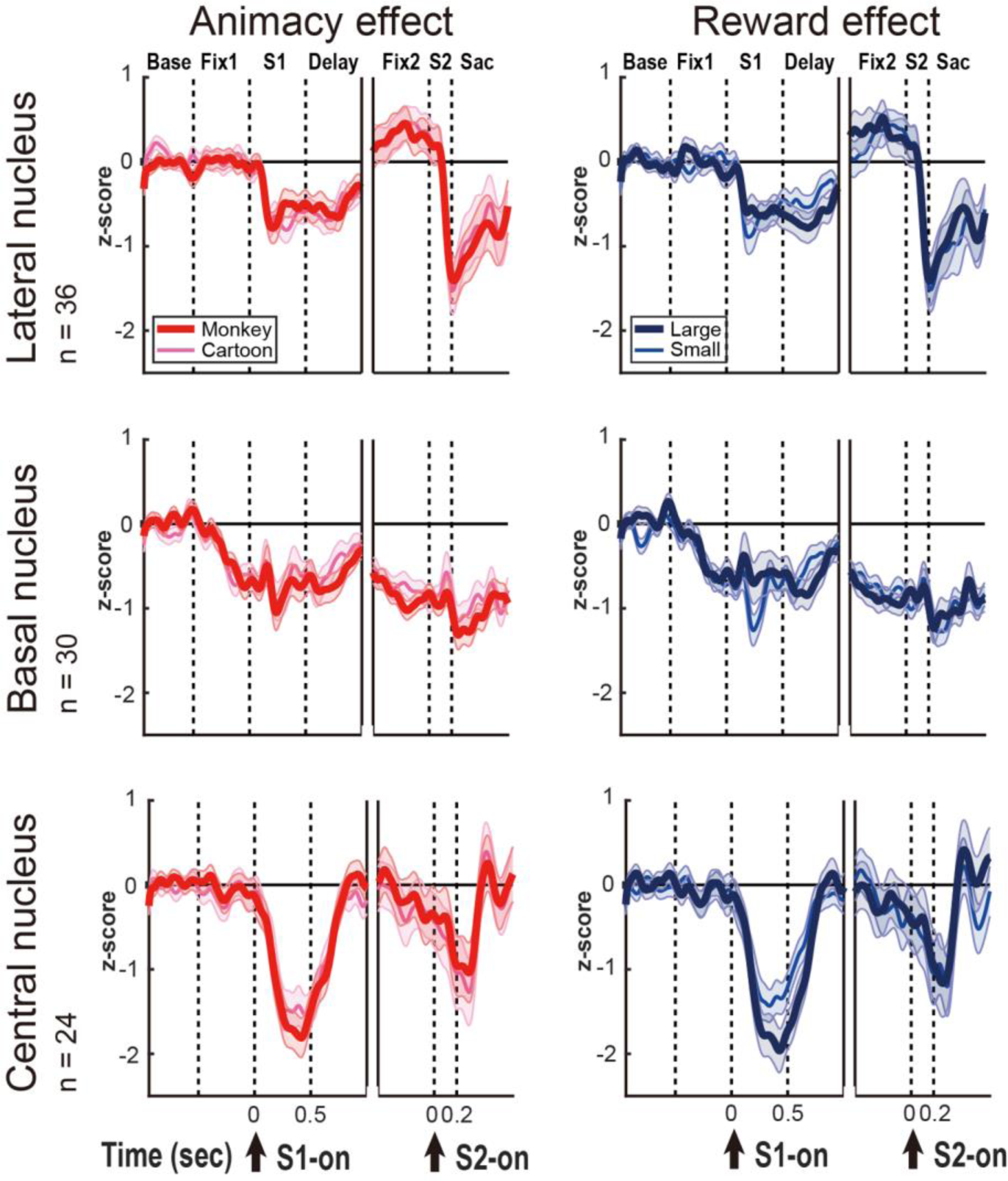
Population inhibitory neuronal responses to S1 and S2.

**Supplementary Figure 4-1.**
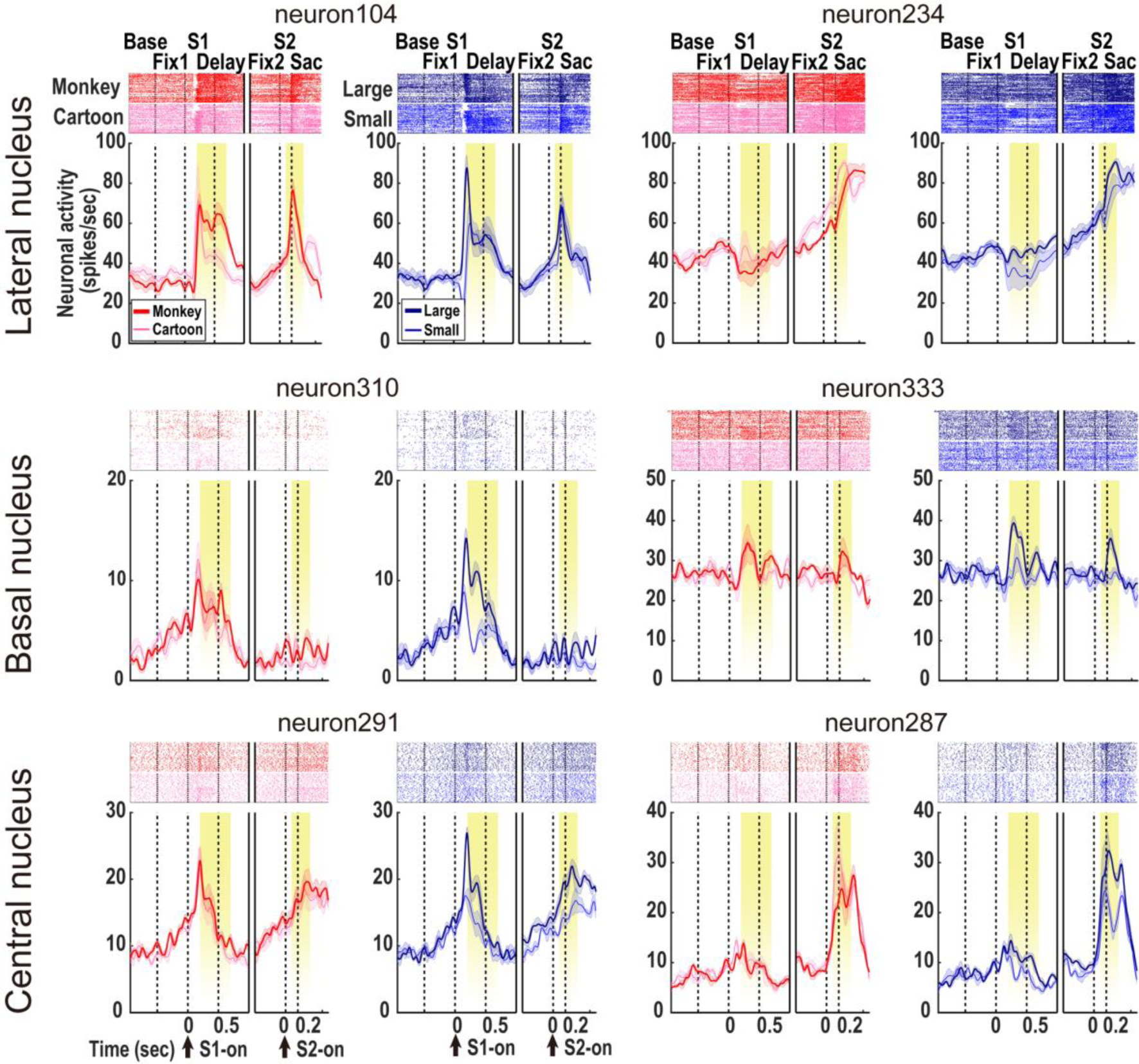
Other examples of excitatory single-neuron responses to the S1 and S2 in the three amygdala nuclei. We compared the neuronal activity elicited by the monkey versus cartoon face stimuli (animacy effect) or the face stimuli associated with the large versus small reward (reward effect). The yellow area in each diagram indicates the periods considered in the analyses.

